# Contrasted hybridization patterns between two local populations of European wildcats in France

**DOI:** 10.1101/342576

**Authors:** Beugin Marie-Pauline, Salvador Olivier, Leblanc Guillaume, Queney Guillaume, Natoli Eugenia, Pontier Dominique

**Author notes:** Corresponding authors: BEUGIN Marie-Pauline, 0786501898, PONTIER Dominique.

## Abstract

The European wildcat (*Felis silvestris silvestris*) is threatened across the totality of its area of distribution by hybridization with the domestic cat *F.s. catus*. The underlying ecological processes promoting hybridization, remain largely unknown. In France, wildcats are mainly present in the North-East but signs of their presence in the Pyrenees have been recently provided. However, no studies have been carried out in the French Pyrenees to assess the genetic status of wildcats. We have compared a local population of wildcats living in a continuous habitat in the French Pyrenees and a local population of wildcats living in a fragmented habitat in Northeastern France to evaluate how habitat fragmentation influence the population structure of European wildcats. Close kin were not found in the same geographic location contrary to what was observed for females in the Northeastern wildcat population. Furthermore, there was no evidence of hybridization in the Pyrenean wildcats and only one domestic cat raised suspicions while hybridization was categorically detected in northeastern France. The two wildcat populations were significantly differentiated (*F_st_* = 0.08) and the genetic diversity of the Pyrenean wildcats was lower than that of other wildcat populations in France and in Europe. Taken together, these results suggest that habitat fragmentation, and in particular the absence of agricultural fields, may play an important role in lowering the probability of hybridization by reducing the likelihood of contact with domestic cats. Moreover, our results suggest that the French Pyrenean wildcat populations is isolated and may be threatened by a lack of genetic diversity.

## INTRODUCTION

Hybridization is especially common between subspecies, due to incomplete reproductive isolation and therefore a higher likelihood of successful interbreeding (1–3). This is the case for the European wildcat *Felis silvestris silvestris*, a medium-sized carnivore widely spread in Europe (4), which is highly threatened over its entire distribution area by its closely related domestic counterparts *F. s. catus* (5). Interbreeding between wildcats and domestic cats may lead to introgressive hybridization, followed by disruption of local genetic adaptations, and then to a loss of the European wildcat genetic integrity, and even to the extinction of the sub-species (6).

Studies across the area of distribution of the European wildcat have shown that there is a high degree of variability in the extent of admixture with domestic cats. High levels (up to 45%) of hybridization have been reported in Hungary and Scotland (3,7–10), while low levels (between 0 and 2%) of interbreeding with domestic cats have been shown in Germany, Italy, and Portugal (9–12). The direction of the gene flow also varied, some studies reporting a gene flow from domestic cats to wildcats (13,14) while others showed the opposite with a detected flow from wildcats to domestic cats (15). Such high degree of heterogeneity in hybridization modalities and subsequent introgression may reflect different environmental conditions (e.g., habitat fragmentation, urban pressure). Characterizing the patterns and processes of hybridization in nature is crucial to the introduction of measures designed to prevent hybridization and then plan efficient conservation guidelines for European wildcats.

The different underlying processes leading to hybridization have been under investigated in the past, since this requires to focusing on the interacting populations of wildcats and domestic cats at a local scale. To our knowledge, only one study combining genetics and radio-tracking of wildcats (16) has been conducted at a local scale in an area of ancient sympatry in Northeastern France. This study was conducted in a fragmented landscape, where forests, crops and villages alternate. A spatial sexual segregation was observed, with females living mostly inside the forest with an access to crops, and males remaining at the edge of the forest. The localization of male wildcats was proposed to be a factor promoting hybridization between them and female domestic cats. Furthermore, the home-ranges of related females were spatially close - even overlapping - suggesting that prey might be abundant enough in this habitat to result in a greater tolerance for overlap between females and their relatives (16). However, this single study may not have accurately captured all the diversity in the hybridization processes and more local studies are urgently needed to deepen our understanding of the factors promoting hybridization in the European wildcat.

In this paper, we extent our analyses in the local population of Northeastern France (16) with a comprehensive sampling of domestic cats allowing for the detection of hybrids both in the wild and the domestic population. In addition, we have conducted a genetic study of wildcats at a local scale within a protected area in the French part of the Pyrenees. The French Pyrenean wildcat population is suspected to be relatively isolated within the species’ distribution range in France and northern Europe (17), and has never been genetically characterized, particularly regarding hybridization. In this area the forest landscape is highly continuous, contrary to the fragmented forests of Northeastern France (18). Here, we have endeavored to assess the genetic status of this wildcat population and the impact of spatial proximity of wildcats to human influences on their genetic admixture by comparing results of relatedness and hybridization pattern with the wildcat population of northeastern France. Given the continuous forest habitat with few interfaces between forests and villages in this area, we may expect hybridization to be rare or inexistent in Pyrenean European wildcats. We did not expect female natal philopatry when food resources are less abundant in the study area due to the absence of agricultural fields.

## MATERIAL AND METHODS

### Study areas and non-invasive sampling

In Northeastern France, the area of study covered approximately 400 km^2^ (16). The landscape is substantially fragmented (18) and consists of an alternation between forests, agricultural fields and permanent grass with elevations ranging from 250-400m. A total of sixteen villages (30 to 600 inhabitants per village) were in direct proximity with the forest where wildcats were sampled. In the Pyrenees, the Nohèdes Nature Reserve presents elevations ranging from 760-2,459m while the elevation of the Jujols Nature Reserve ranges between 1,100 and 2,172m. The study area covers a total surface of 325 km^2^ of continuous forest (oak, maple, ash, pines, beech). These two nature reserves are in proximity with ten villages (30 to 230 inhabitants per village), and more particularly with the village of Nohèdes, Urbanya, Conat and Jujols.

In Northeastern France, domestic cats (N = 371) and wildcats (N = 32) were captured, and blood samples were collected for each individual, from November–February of each year between April 2008 and May 2011, using trapping cages containing crushed valerian roots (*Valeriana officinalis*), a common attractant for cats. Trapped individuals were anaesthetized with ketamine chlorohydrate (Imalgène 1000, 15 mg/kg, Merial) and aceprozamine (Vétranquil 5.5 %, 0.5 mg/kg, Ceva). A permanent subcutaneous electronic device (transponder Trovan, AEG & Telefunken Electronic, UK) was injected in each cat to aid subsequent identification. In addition, ten wildcat samples were obtained on road-killed individuals. The fieldwork has been conducted by qualified people according to current French legislation. Accreditation has been granted to the UMR-CNRS5558 (accreditation number 692660703) for the program. In the Pyrenees, fresh feces of wildcats were collected opportunistically from 2010 to 2016 in the nature reserves of Jujols and Nohèdes in the eastern part of the French Pyrenees (Figure 1). Experienced field agents from the nature reserves have collected evidence for additional occurrences of European wildcats since 1993 based on camera-trapping surveys, direct observations or feces, which has allowed us to associate the sampling of fresh feces to the overall presence of the European wildcat in the study area. For domestic cats, hairs were sampled in 2010 and 2017 in the villages of Nohèdes (N = 20), Conat (N=4), and Serdinya (N = 3), located on the edge of the reserves (see Figure 1). Only individuals born in the villages, sterilized or not, were included in our sampling. A total of 71 feces and 27 hair samples were thus collected. Feces were sampled in individual plastic bags and kept at −20°C; and hairs in dry closed envelopes.

**Figure 1:**
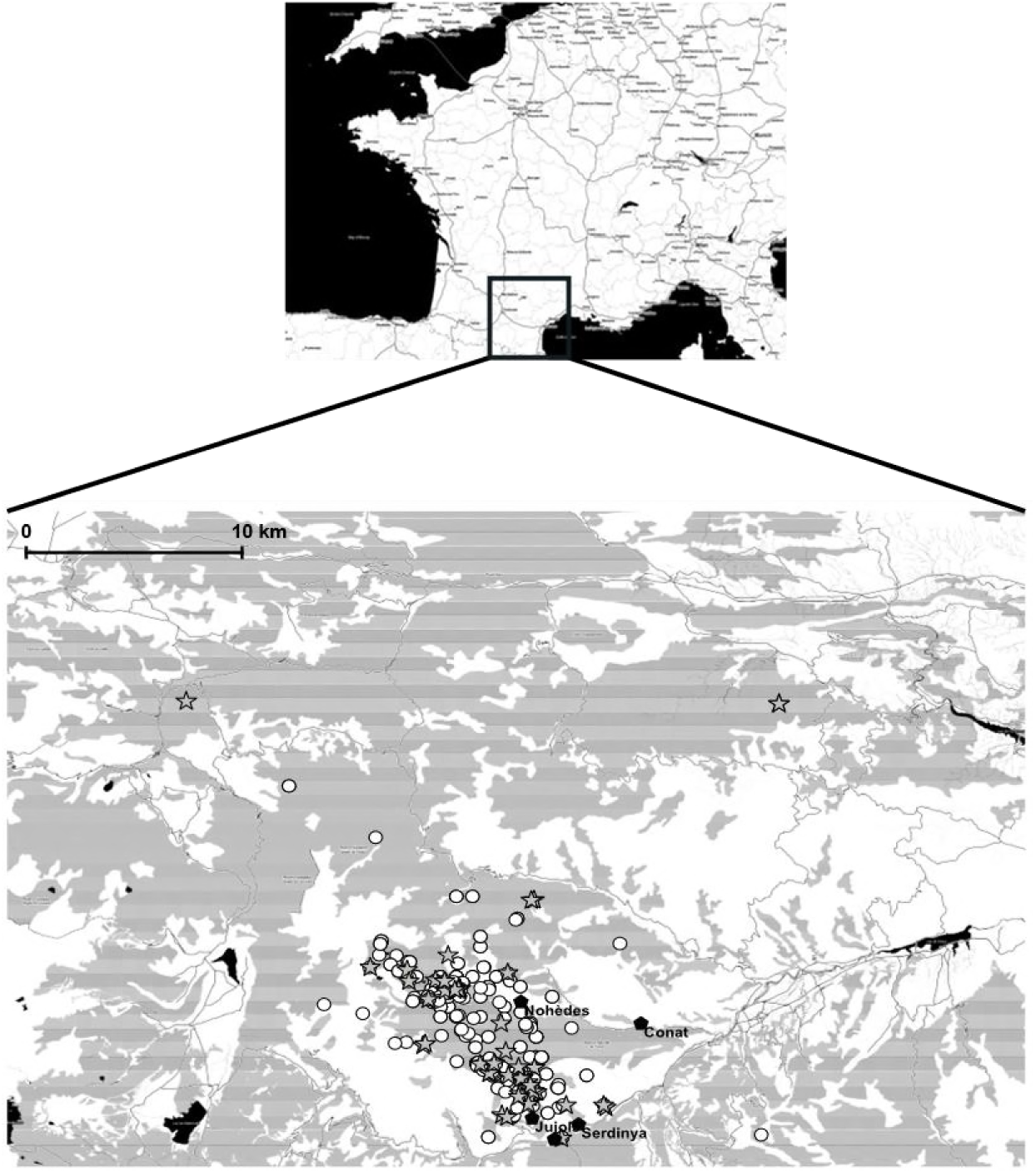
Localization of the study area in the Pyrenees. The circles represent all the locations where feces, camera-trapping or direct observations attested for the presence of the European wildcat. The stars correspond to the locations of the fresh feces. The villages where domestic cats were sampled are indicated in black and are located at the edge of the natural reserves of Jujols and Nohèdes.

### DNA extraction and microsatellite genotyping

Northeastern domestic cats were genotyped using 22 autosomal microsatellites, Pyrenean cats and Northeastern wildcats were genotyped using 31 autosomal microsatellites. All cats were genotyped with a marker of sex. DNA extraction was performed using a purification column kit (Nucleospin 96 Tissue kit, Macherey-Nagel) following the manufacturer protocol. PCR reactions were performed step-by-step following a unidirectional workflow starting in a clear room with positive air pressure where sensitive reagents, enzymes and primers, were prepared. DNA and reagents were then assembled in a pre-PCR room. PCR amplifications were made in 96-well microplates in a post-PCR area with negative air pressure. The PCR reaction occurred in a final volume of 10µl that contained 5µl of Mastermix Taq polymerase (Type-it, QIAGEN), 1.35µl of primer pairs at a final concentration between 0.08 and 0.6µM, and 30ng of DNA. Each pair of primers was coupled with a fluorescent dye. The reaction started with a denaturation step at 95°C for five minutes. This step was followed by thirty PCR cycles (denaturation step = 95°C, 30s; annealing step = 55.9°C, 90s; elongation step = 72°C, 30s) and a final elongation step at 60°C during 30 minutes. PCR products were resolved on a capillary sequencer ABI PRISM 3130 XL (Applied Biosystem) under denaturing conditions (formamide) and an internal size marker in one migration for each multiplex. All these steps were performed using filtered tips. Finally, the electrophoregrams were analyzed using GENEMAPPER 4.1 (Applied Biosystem/Life Technologies) twice independently. Ambiguous loci were classed as missing data.

Blood samples (northeastern France samples) were genotyped only once. For non-invasive samples, a first genotyping was conducting on all samples. Those for which none or only a few loci were amplified were discarded, the other samples were genotyped one to three additional times. A consensus genotype was then built to obtain a genotype per sample. A quality index (referred to as QI thereafter, 19) was calculated in order to assess the quality of the samples. For each locus and each repeat, the genotype obtained is compared to the consensus genotype built over all repeats. A repeat where the consensus genotype is obtained is rated with QI equal to 1, a repeat with a homozygote genotype while the consensus genotype is heterozygote will have a QI equal to 0.5, and a missing data will correspond to QI equal to 0. The QI are then averaged over all repeats for each locus, and then over all loci for an individual to obtain a QI per sample. Only individuals presenting a QI superior than 0.6 were included in the subsequent analyses.

### Consensus genotypes and population genetics analyses

Consensus genotypes were built as follows. Two genotypes were considered to represent the same individual when (1) they were identical, (2) they only differed by missing data and these missing data did not represent more than ten microsatellite markers, (3) they only differed by missing data below the threshold of ten markers and a single difference that could be explained by allelic dropout. Deviations of loci from both Hardy-Weinberg equilibrium (HWE) and linkage equilibrium were both tested using FSTAT v 2.9.3.2. (20) with a Bonferroni correction and a 5% risk for all populations. Loci showing a departure from HWE were discarded from the analysis. For each locus, the frequency of null alleles was assessed following Brookfield’s (21) method, and its impact on a possible deviance from Hardy-Weinberg equilibrium tested using binomial tests according to De Mêeus et al. (22). All loci exhibiting significant evidence of null alleles causing HWE were discarded from further analyses. The software FSTAT v2.9.3.2. was also used for estimating Weir and Cockerham’s *F_ST_* between wildcat and domestic cat populations from the Pyrenees and from Northeastern France as well as allelic richness. Expected (*H_E_*) and observed (*H_0_*) heterozygosities were calculated using GenALEx 6.501 (23). Finally, a discriminant analysis of principal components (DAPC, 24) was used in order to visualize the differentiation between domestic and European wildcat populations from northeastern France and the Pyrenees.

### Spatial structure and relatedness

The 52 fresh fecal samples for which the sampling location was recorded were displayed on a map using QGIS v2.8.1., together with the other indices of presence of European wildcats (feces and photo trapping). The program ML-Relate (25) was used to calculate pairwise relatedness between all wildcat individuals. Using a linear model, we tested whether sex or relatedness were a significant predictor of the pairwise geographical distance between individuals. Geographical distances between individuals were calculated with QGIS v2.8.1. We considered the mean pairwise distance between samplings for individuals sampled several times. Statistical analysis was performed in R 3.3.3.

### Admixture analysis

The Bayesian clustering method implemented in STRUCTURE v2.3.4 (26,27) was used to identify wildcats, domestic cats and possible hybrids by applying the *admixture model* with correlated allele frequencies. The optimal number of clusters *K* was determined using the method described by Evanno et al. (28). STRUCTURE was used to assess membership proportions (*q_i_*) to the inferred *K* clusters, which correspond to the proportion of each individual’s multilocus genotype belonging to each of the inferred *K* clusters. The threshold level to differentiate wildcats and domestic cats from hybrids was determined by selecting individuals which were believed to be representative of the parental populations using the iterative algorithm described in Beugin et al. (in preparation). This algorithm consists of the repetition of a simulation-selection process allowing for the building of a representative pool of parents (representative regarding the differentiation between parental population and their genetic diversity). At each step, individuals from different hybrid classes (parental classes, F1, F2, first generation backcrosses) were simulated using the function *hybridize* from the package *adegenet* (29). At initialization, these simulations were carried out based on the 5% top-ranked (based on *q-*values) individuals from each cluster. These simulated individuals were then analyzed with STRUCTURE using no prior information in order to get individual *q-*values. All the individuals characterized by a lower bound of their 90% credibility interval higher than the 1% quantile of the distribution of *q-*values of the parental simulated individuals were then integrated into the representative parental pool and used to carry on the simulations at the next iteration. This simulation-selection process was then repeated until the number of individuals integrated in the representative parental pool stabilized. This way, all individuals that can be considered as a parental individual are used to define the threshold that allows hybrids to be distinguished from parents.

In order to carry out such categorization, we simulated 200 individuals (domestic cats, wildcats, F1, F2, F1 x domestic and F1 x wild) using the function *hybridize* from the R package *adegenet* (29) based on the pool of representative individuals built and ran STRUCTURE ten times in order to determine four thresholds corresponding to the lowest *q-*values reached by parental individuals on one hand (thresholds TP1 and TP2) and the highest *q-*values reached by hybrid individuals in the second hand (thresholds TH1 and TH2). This defined the borders of the parental zone, the hybrid zone and the grey zone. All STRUCTURE analyses (iterative algorithm and threshold determination) were run for a burn-in period of 100,000 and MCMC length of 100,000 iterations according to Gilbert et al. (30) and following graphical verifications of the convergence of the algorithm based on the parameter alpha. We ran the function *snapclust* (31) from the *adegenet* package in order to determine the category of detected hybrids. Given the number of microsatellites and the expected order of magnitude of the differentiation between domestic cats and wildcats (*F_st_* = 0.11 to 0.20 – ref 7,12,15,16,32) we considered only F1 hybrids and first-generation backcrosses (F1 x Domestic cat and F1 x Wildcat) in the *snapclust* analysis. Additionally, the direction of the gene flow between wild and domestic cats was assessed by estimating the rate of migration per generation with the computer program BAYESASS 3.0.3. (33) with a MCMC chain of 5,000,000 after a burn-in period of 1,000,000 with a sampling interval of 2,000. All other parameters were left to default.

## RESULTS

### Amplification success and characterization the Pyrenean population

Feces and/or direct observations of European wildcats have been reported all over the nature reserves of Jujols and Nohèdes up to an elevation of 2430m. Samples genotyped were collected over the entire area where signs of the presence of the European wildcat had been reported (see Figure 1). Forty-four out of the 71 fresh feces collected, and 22 out of the 27 hair samples collected showed a QI superior than 0.6 (mean = 0.84, sd = 0.12, see supplementary material, Figure S1). On average, 74.1% (sd = 0.31) of the loci were successfully amplified using feces while 81.5% (sd = 0.32) of the loci were successfully amplified on average from hairs.

We identified 39 unique genotypes including 21 domestic cats (15 females and 6 males) and 18 European wildcats (10 females and 8 males). Six (3 females and 3 males) out of the 18 European wildcats were sampled several times (from two to 6 times). Feces from the same individual were found at a maximal distance of 5.3 km for a male, and 3 km for a female. Different feces from the same individual were found within 540 m of elevation for one male, whereas the other individuals were sampled within 100-200 m of elevation. These recaptures did not allow us to establish home-ranges due to the lack of locations per individual. We detected wildcats genetically confirmed as such in elevations up to 2250 m.

### Genetic diversity and kinship pattern

We did not detect any linkage disequilibrium in any of the populations. On the contrary, the northeastern population of domestic cats showed signs of Hardy-Weinberg disequilibrium for six loci (Fca8, Fca45, Fca96, Fca229, Fca453). The presence of null alleles did not explain significantly these deviations.

Domestic cats showed a higher allelic richness than European wildcats both in the Pyrenees (mean = 6.55, sd = 1.34 for domestic cats, mean = 4.84, sd = 1.46 for wildcats, Table 1) and in Northeastern France (mean = 7.5, sd = 2.94 over 31 domestic cats, mean = 6.74, sd = 1.95 for wildcats). Wildcats from Northeastern France presented more alleles per locus on average than their Pyrenean conspecifics. Average values of heterozygosity were slightly higher in the Pyrenean domestic cats (*H_0_* = 0.71 *H_E_* = 0.73) than in European wildcats (*H_0_* = 0.67; *H_E_* = 0.62), and only expected heterozygosity was higher in Northeastern France. Overall, heterozygosities between domestic cats and wildcats were very similar.

**Table 1:** Mean number of alleles (Na), observed heterozygosity (Ho) and expected heterozygosity (He) per locus for each population of wildcats and domestic cats: Pyrenean wildcats (PO wildcats), Northeastern wildcats (NE wildcats), Pyrenean domestic cats (PO domestic cats) and northeastern domestic cats (NE domestic cats). An additional column gives estimates of genetic diversity for a subset of northeastern domestic cats that could be genotyped over 31 microsatellites. Loci that were not genotyped or for which genotyping failed are indicated with ‘-‘.

|  | PO WILDCATS (N = 19) |  |  | NE WILDCATS (N = 42) |  |  | PO DOMESTIC CATS (N = 21) |  |  | NE DOMESTIC CATS (N = 371) |  |  | NE DOMESIC CATS (N = 31) |  |  |
| --- | --- | --- | --- | --- | --- | --- | --- | --- | --- | --- | --- | --- | --- | --- | --- |
| MARKER | Na | Ho | He | Na | Ho | He | Na | Ho | He | Na | Ho | He | Na | Ho | He |
| F37 | 3 | 0.25 | 0.40 | 5 | 0.19 | 0.33 | 8 | 0.92 | 0.84 | - | - | - | - | - | - |
| FCA8 | 7 | 0.79 | 0.75 | 8 | 0.89 | 0.81 | 7 | 0.85 | 0.80 | - | - | - | 9 | 0.71 | 0.87 |
| FCA23 | 3 | 0.74 | 0.61 | 4 | 0.60 | 0.62 | 6 | 0.54 | 0.75 | - | - | - | - | - | - |
| FCA24 | 5 | 0.46 | 0.70 | 9 | 0.80 | 0.80 | 6 | 0.92 | 0.79 | - | - | - | - | - | - |
| FCA31 | 7 | 0.67 | 0.67 | 8 | 0.68 | 0.65 | 8 | 0.62 | 0.80 | - | - | - | 9 | 0.70 | 0.84 |
| FCA43 | 6 | 0.89 | 0.80 | 6 | 0.76 | 0.73 | 7 | 0.57 | 0.59 | - | - | - | 5 | 0.74 | 0.66 |
| FCA45 | 5 | 0.68 | 0.70 | 8 | 0.73 | 0.76 | 8 | 0.71 | 0.80 | 16 | 0.76 | 0.86 | 12 | 0.77 | 0.80 |
| FCA58 | 4 | 0.37 | 0.32 | 7 | 0.45 | 0.50 | 6 | 0.90 | 0.79 | 11 | 0.62 | 0.70 | 7 | 0.55 | 0.59 |
| FCA77 | 6 | 0.74 | 0.65 | 8 | 0.83 | 0.82 | 5 | 0.52 | 0.47 | - | - | - | 6 | 0.61 | 0.63 |
| FCA78 | 5 | 0.53 | 0.52 | 7 | 0.62 | 0.75 | 7 | 0.50 | 0.76 | - | - | - | 5 | 0.80 | 0.74 |
| FCA85 | 2 | 0.32 | 0.27 | 4 | 0.64 | 0.61 | 5 | 0.71 | 0.64 | - | - | - | 5 | 0.40 | 0.63 |
| FCA96 | 7 | 1.00 | 0.79 | 10 | 0.90 | 0.85 | 6 | 0.45 | 0.45 | 15 | 0.37 | 0.42 | 11 | 0.55 | 0.60 |
| FCA124 | 7 | 0.89 | 0.83 | 9 | 0.79 | 0.81 | 7 | 0.86 | 0.80 | 10 | 0.76 | 0.78 | 8 | 0.84 | 0.82 |
| FCA126 | 6 | 0.63 | 0.73 | 8 | 0.66 | 0.74 | 10 | 0.78 | 0.87 | 10 | 0.53 | 0.58 | 6 | 0.35 | 0.41 |
| FCA547 | 5 | 0.79 | 0.72 | 6 | 0.71 | 0.70 | 7 | 0.67 | 0.78 | - | - | - | 9 | 0.77 | 0.80 |
| FCA577 | 4 | 0.67 | 0.66 | 7 | 0.56 | 0.51 | 7 | 0.69 | 0.77 | 14 | 0.74 | 0.79 | 5 | 0.63 | 0.72 |
| FCA668 | 4 | 0.83 | 0.69 | 7 | 0.73 | 0.68 | 6 | 0.71 | 0.71 | 13 | 0.82 | 0.86 | 10 | 0.81 | 0.83 |
| FCA675 | 6 | 0.74 | 0.79 | 8 | 0.74 | 0.80 | 10 | 0.90 | 0.83 | 14 | 0.76 | 0.79 | 10 | 0.87 | 0.76 |
| FCA26 | 6 | 0.74 | 0.68 | 9 | 0.93 | 0.84 | 6 | 0.67 | 0.74 | 17 | 0.79 | 0.83 | 10 | 0.90 | 0.85 |
| FCA69 | 5 | 1.00 | 0.76 | 7 | 0.69 | 0.72 | 7 | 0.81 | 0.80 | 14 | 0.79 | 0.82 | 7 | 0.65 | 0.78 |
| FCA75 | 5 | 0.78 | 0.72 | 9 | 0.81 | 0.82 | 5 | 0.71 | 0.72 | 11 | 0.71 | 0.74 | 7 | 0.74 | 0.72 |
| FCA105 | 7 | 0.84 | 0.80 | 9 | 0.74 | 0.75 | 7 | 0.67 | 0.66 | 17 | 0.77 | 0.83 | 9 | 0.74 | 0.75 |
| FCA149 | 4 | 0.56 | 0.57 | 4 | 0.71 | 0.71 | 5 | 0.81 | 0.72 | 7 | 0.81 | 0.78 | 6 | 0.84 | 0.78 |
| FCA201 | 4 | 0.74 | 0.54 | 7 | 0.76 | 0.74 | 8 | 0.86 | 0.83 | 14 | 0.78 | 0.80 | 9 | 0.77 | 0.80 |
| FCA220 | 5 | 0.75 | 0.73 | 7 | 0.83 | 0.84 | 6 | 0.71 | 0.64 | 10 | 0.48 | 0.51 | 8 | 0.53 | 0.49 |
| FCA229 | 4 | 0.47 | 0.51 | 8 | 0.71 | 0.66 | 5 | 0.62 | 0.62 | 11 | 0.66 | 0.76 | 6 | 0.59 | 0.73 |
| FCA293 | 3 | 0.83 | 0.66 | 4 | 0.71 | 0.71 | 6 | 0.67 | 0.79 | 8 | 0.75 | 0.76 | 6 | 0.68 | 0.71 |
| FCA310 | 2 | 0.05 | 0.05 | 3 | 0.17 | 0.23 | 6 | 0.57 | 0.64 | 8 | 0.65 | 0.70 | 6 | 0.52 | 0.68 |
| FCA441 | 5 | 0.71 | 0.53 | 4 | 0.64 | 0.66 | 5 | 0.71 | 0.73 | 9 | 0.69 | 0.68 | 6 | 0.71 | 0.70 |
| FCA453 | 3 | 0.54 | 0.50 | 4 | 0.69 | 0.67 | 6 | 0.71 | 0.68 | 8 | 0.64 | 0.72 | 7 | 0.77 | 0.73 |
| FCA678 | 5 | 0.83 | 0.69 | 5 | 0.69 | 0.64 | 5 | 0.71 | 0.72 | 7 | 0.70 | 0.76 | 6 | 0.58 | 0.75 |
| Mean | 4.84 | 0.67 | 0.62 | 6.74 | 0.69 | 0.69 | 6.55 | 0.71 | 0.73 | 11.62 | 0.69 | 0.74 | 7.50 | 0.68 | 0.72 |
| SD | 1.46 | 0.22 | 0.17 | 1.95 | 0.17 | 0.14 | 1.34 | 0.13 | 0.10 | 6.13 | 0.34 | 0.36 | 2.94 | 0.24 | 0.24 |

European wildcats and domestic cats were significantly differentiated with a *F_ST_* value of 0.18 in the Pyrenees, 0.15 in northeastern France. A significant differentiation between Pyrenean wildcats and Northeastern domestic cats (*F_ST_* = 0.19), and wildcats (*F_ST_* = 0.0835), was also found, while Northeastern domestic cats were significantly differentiated from Pyrenean domestic cats (*F_ST_* =0.0414). The DAPC scatter-plot (Figure 2) confirmed the sharp distinction between the two European wildcat populations.

**Figure 2:** Scatter plot of the discriminant analysis of principal components (DAPC) integrated all Pyrenean domestic cats and wildcats, northeastern wildcats, and 31 domestic cats that could be genotyped over the 31 microsatellites. The first PC describes 81% of the genetic diversity and the second axis 14%.

Females from the Pyrenean wildcat population showed an *F_is_* value close to zero (*F_is_* = 0.00028), while males showed a heterozygote excess (*F_is_* = −0.09). Wildcat females also presented a higher *F_is_* than males in northeastern France but with females being inbred (*F_is_* = 0.04) and males at equilibrium (*F_is_* = −0.01). Individuals from the Pyrenean wildcat population were poorly related and, in both sexes, related wildcat individuals were not sampled significantly closer together compared to unrelated individuals in the Pyrenees according to the linear mixed model (Supplementary Material, Figure S3). On the contrary, in Northeastern France, related females were captured significantly closer together than unrelated females (F = 6.88, df = 1, p = 0.0095; Supplementary Material, Figure S3).

### Admixture analysis

Evanno’s method showed that *K*=2 best described our data in both populations, one cluster corresponding to the domestic cats, the other corresponding to the European wildcats (Figure 3). The iterative algorithm allowed us to define hybrids as individuals presenting a mean probability of assignment (conservative approach) or a lower bound of credibility interval (relaxed approach) below 0.79 for domestic cats, and below 0.83 for European wildcats in the Pyrenees; below 0.82 for domestic cats and below 0.89 for European wildcats in Northeastern France.

**Figure 3:**
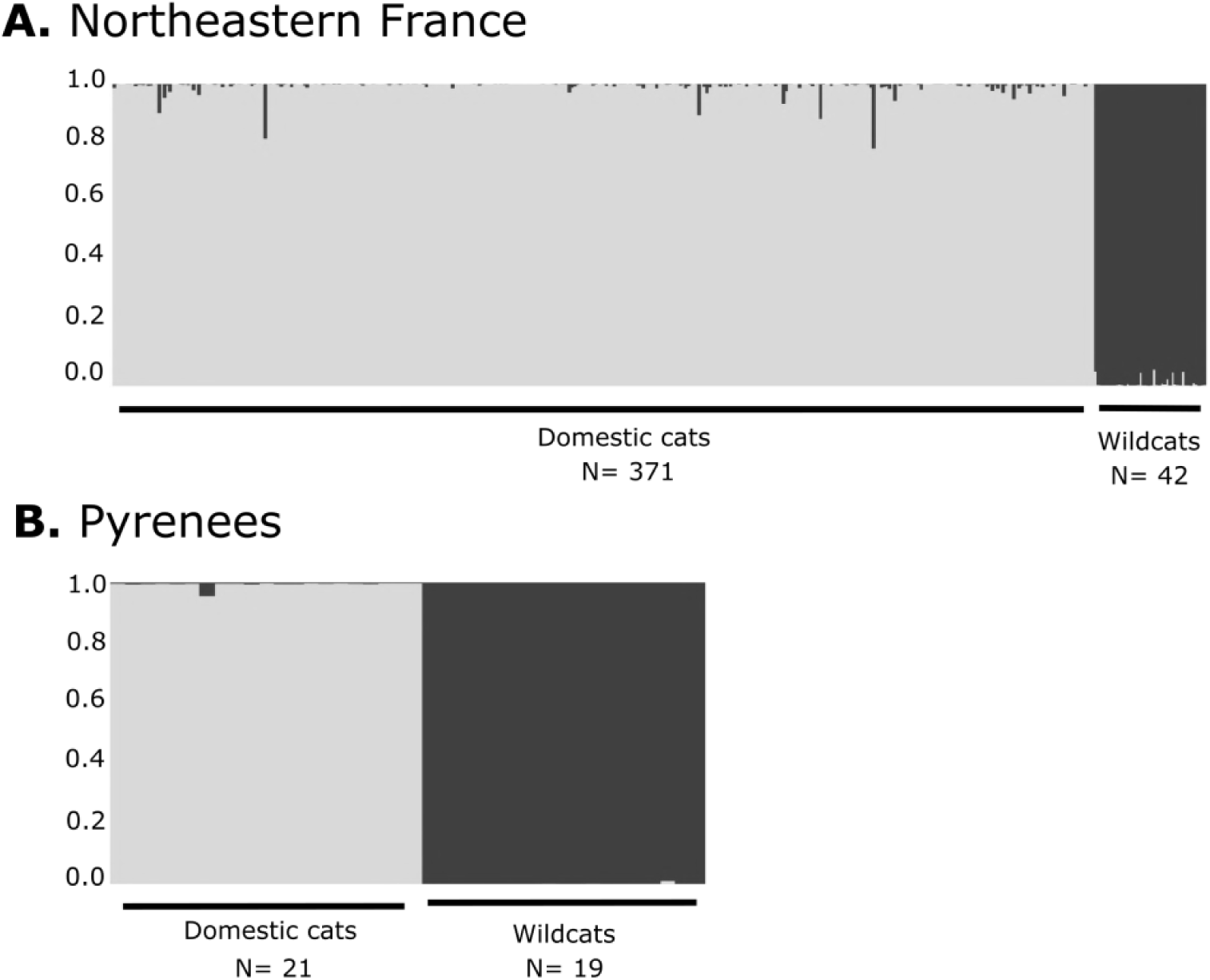
Results from the STRUCTURE analysis in northeastern France (A) and in the Pyrenees (B). Each cat genotype is represented by a vertical bar split into *K*=2 colored sections, according to its relative assignment to the genetic cluster. The proportion of the bar in a given color represents the assignment probability of the individual for the corresponding cluster.

Using the conservative approach, we did not detect any hybrid and *snapclust* confirmed the absence of hybrids in the Pyrenees, while the relaxed approach detected one of the domestic cats sampled in Nohèdes as being hybrid and this individual detected as being hybrid by the relaxed approach was substantially assigned to the first-generation backcross category by *snapclust* (Supplementary material, Figure S4). The absence of gene flow from domestic cats to wildcats (m = 0.0148 – CI95: 0-0.042) as well as from wildcats to domestic cats (m = 0.0160 – CI95: 0-0.046) was consistent with the absence of hybridization between the two sub- species. On the contrary, in Northeastern France, hybrids were systematically detected, at least in the domestic cat population. Thus, using the conservative approach, two hybrids were detected in domestic cats and none in the European wildcats while sixteen and six hybrids were detected in the domestic cats and European wildcats respectively using the relaxed approach. All the individuals detected as hybrids with the relaxed approach were substantially assigned to the first-generation backcross category by *snapclust* (Supplementary material, Figure S5). No significant gene flow was detected from domestic cats to wildcats (m = 0.0009 – CI95: 0-0.0034) while a low but significant gene flow was detected from European wildcats to domestic cats (m = 0.0077 – CI95: 0.00019-0.028).

## DISCUSSION

Our study has confirmed the presence of wildcats in the French Pyrenees within a large area of the nature reserves of Nohèdes and Jujols up to 2,430 m. Moreover, our study has provided the first genetic characterization of a local population of the French Pyrenean wildcats, despite their presence being acknowledged since 1993 (Observations collected by trained personnel of the nature reserve; ref 17).

The primary interesting result is that males as well as females in close proximity are not kin related suggesting that both males and females disperse in this continuous forest landscape, i.e., related females did not tend to remain in the same area contrary to the wildcat population of northeastern France. The dispersal pattern may directly reflect the level of food resource availability. In fragmented environments as observed in northeastern France, with forest alternating with field crops, large areas rich in resources are available for wildcats (34,35). With carnivores, food distribution has been suggested to be the major determinant of species spatial distribution (36). The importance of resource distribution on the spacing pattern of wildcat females has already been proposed (37,38) and was supported by the study in northeastern France (16). Thus, although the European wildcat is acknowledged to live solitarily (39,40), its dispersal pattern may show more variability than has been described up to now.

The second important result is that no hybrid was categorically found neither in the Pyrenean wildcats nor in the Pyrenean domestic cat population. The absence of hybrids in the Pyrenean population contrasts with the situation in northeastern France, where two hybrids were categorically identified among domestic cats and six wildcats out of 42 showed signs of hybridization. The gene flow from wildcats to domestic cats found in this populations is consistent with the one reported by Hertwig et al. (15) in eastern Germany, in a fragmented area notably marked by extensive cultivated landscapes and similar pattern of females spatial distribution (41).

Compared with estimations in the same area, the rate of hybridization we found in northeastern France is surprisingly low (13.8% against 25% on average, 17,42–44). A possible explanation for this difference may be linked to different sampling strategies. While our sampling is local, focused on one population of living wildcats and on neighboring populations of living domestic cats, previous studies in France relied upon opportunistic sampling of road-killed animals, over a much larger area. Germain *et al.* (42,43) showed that hybrids tend to live in intermediary environments, between forests and villages. This would expose them to road mortality more often than wildcats living in the forest or domestic cats in the villages and thus, sampling schemes based on road-killed animals may be biased towards a higher proportion of hybrids.

The absence of hybrids in the continuous Pyrenean forest landscape may imply that the absence of crops frequented by both wild and domestic cats for hunting purposes may impede encounters between the two subspecies as domestic cats, even feral domestic cats, do not enter the forest environment (during the last seven years no pictures of domestic cats have been taken by camera trapping). Unfortunately, the sample size of this study was limited and more extensive studies will be required to confirm the absence of hybridization in this environmental context.

Finally, the Pyrenean wildcats showed values of genetic diversity lower than other wildcat or domestic cat populations in France and in Europe (4.84 alleles per locus on average in the Pyrenean wildcat population while between 3 and 11.8 can be found in the literature with rare populations below 6; ref 13,17,32,44,45), suggesting that wildcats in the Pyrenees may be threatened by a lack of genetic diversity. Not surprisingly, the genetic differentiation between Pyreneans domestic cats and their northeastern counterparts was moderate (*F_ST_* =0.04). In contrast, the Pyrenean wildcat population was significantly differentiated from the northeastern wildcats with a higher *F_ST_* value (0.08). This genetic divergence could result from a classic genetic process of isolation by distance (IBD, 46), which supposes continuity in the distribution of the European wildcat on French territory; such a cline has been described in the northeastern wildcat population (17,47). Alternatively, some gap in the distribution of wildcats may exist in France as suggested by O’Brien et al. (4) and Say et al. (17). The lower genetic diversity of the Pyrenean wildcat population is supportive of the isolation of this population and thus of the existence of a fragmented pattern of distribution of the European wildcat across France.

## CONCLUSION

Results in this study have added novel information to the European wildcat population structure in France. They provided further information about the relationship between environmental conditions and hybridization risks in French wildcat populations. Conservation strategies of wildcats should take into account the local habitat such as the existence of a fragmented or continuous forest environment, and the presence of agricultural fields (41). Further investigation should also focus on the spatial distribution of French wildcats - in particular we need to confirm whether the French Pyrenean population is isolated from the main area of wildcats in France but connected to the Spanish Pyrenean wildcat population. Depending upon the answer, the usefulness of wildlife corridors to enhance connectivity between the different wildcat populations should be addressed to ensure the long-term viability of the French Pyrenean wildcat population.

## ACKNOWLEDGEMENTS

We thank everybody from the reserves and villages of Nohèdes, Conat, Jujols and Serdinya who collaborated in providing wildcat and domestic cat samples and sharing their observations. We also thank Robin Buckland for his careful check of English language. MPB, DP are supported by the LabEx ECOFECT (ANR-11-LABX-0048).

## SUPPLEMENTARY MATERIAL

**Figure S1:**
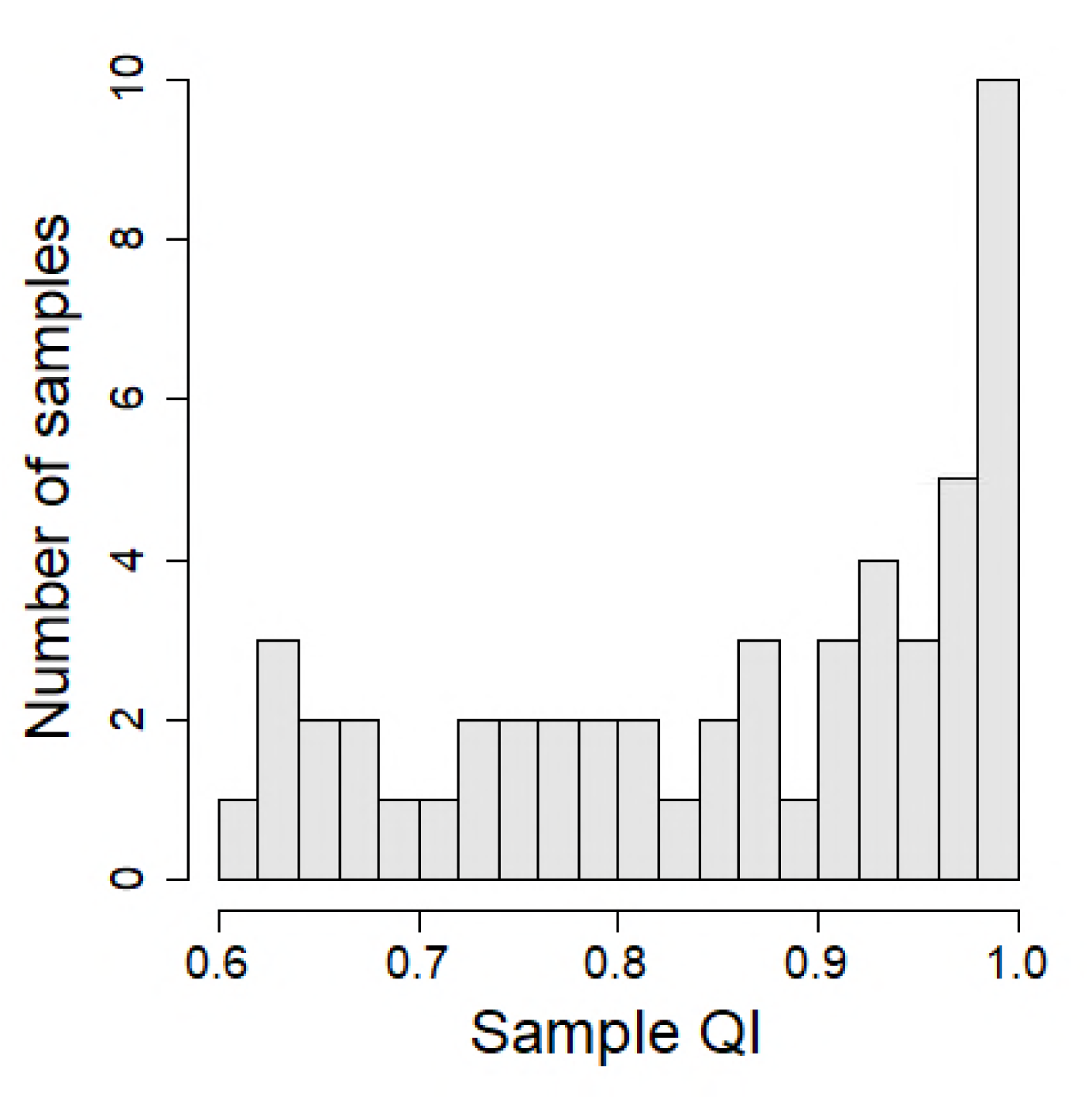
Distribution of the quality indexes of the samples included in the analyses.

**Figure S2:**
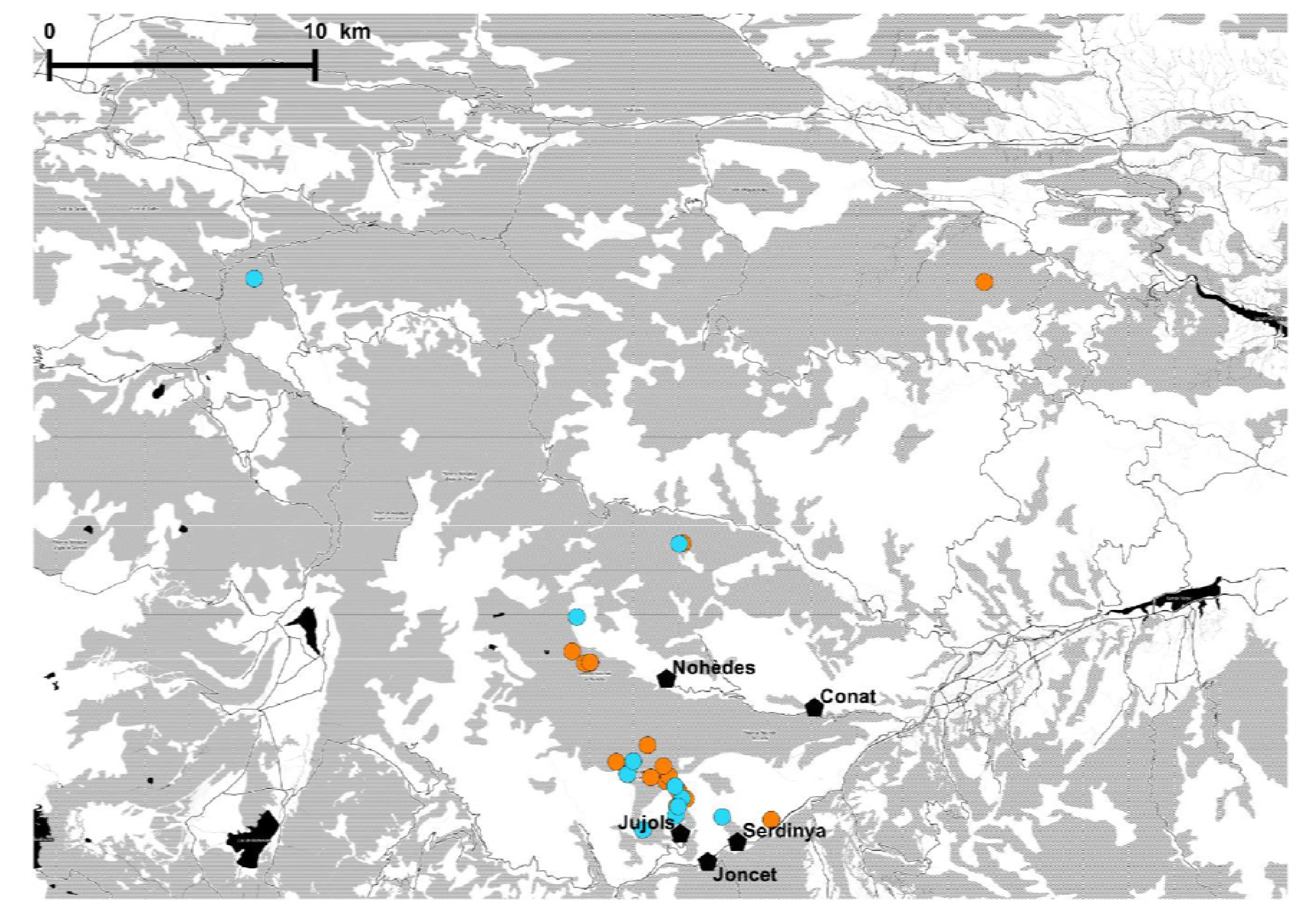
locations of the samples according to the sex of individuals. Females are indicated in orange and males in blue

**Figure S3:**
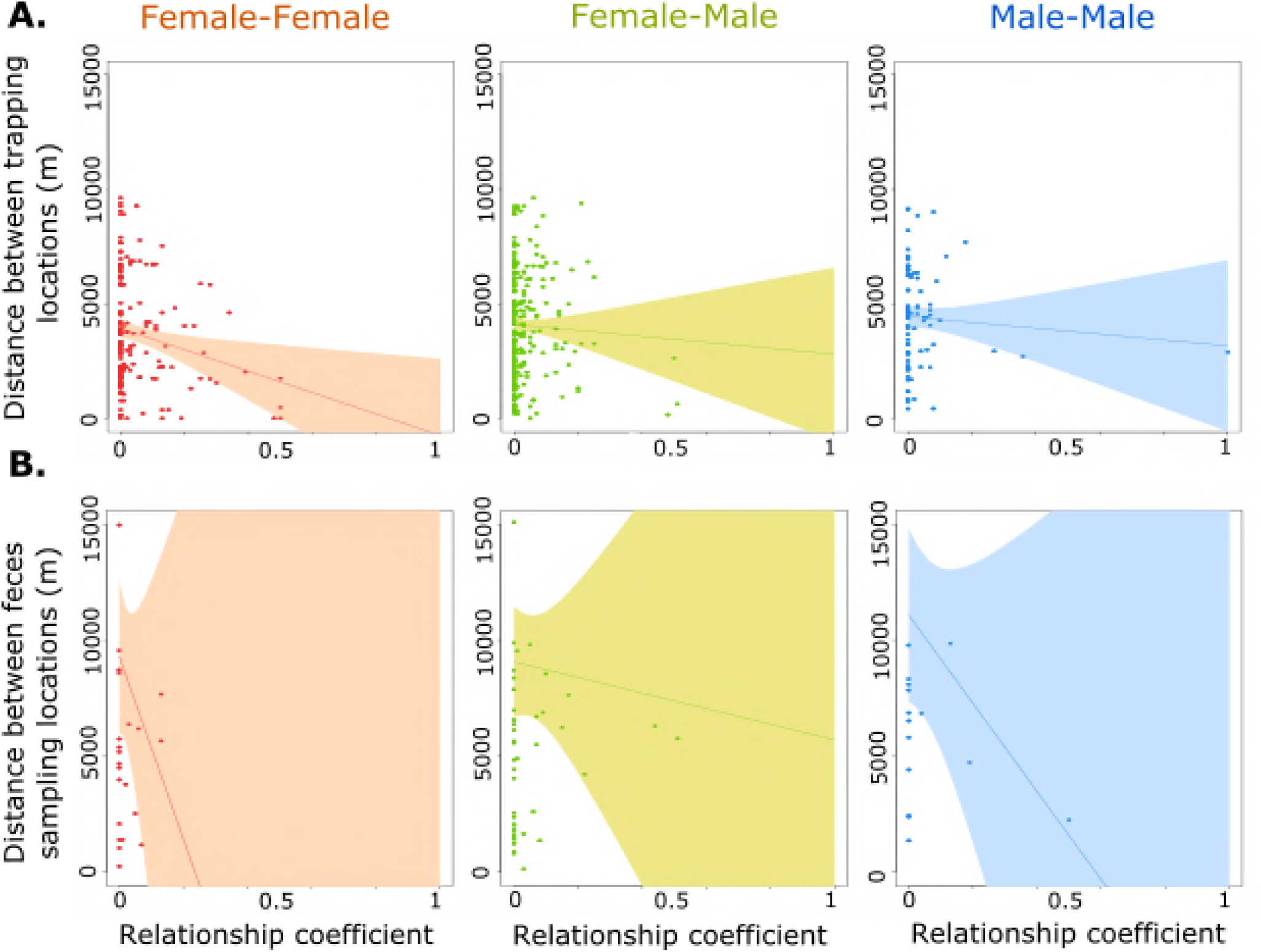
Linear regression and 95% confidence interval (represented using different colors) of the distance between two fresh feces samples according to the coefficient of relationship. A. in northeastern France and B. in the Pyrenees.

**Figure S4:** Results from the *snapclust* analysis for the Pyrenean population. Each vertical bar represents an individual. The colors represent the different hybrid categories included in the analysis: domestic cats in blue, wildcats in green, F1 in purple, F1xdomestic cats in orange, F1xwildcats in brown. The proportion of the bar in a given color represents the probability for the individual to belong to the corresponding category.

**Figure S5:**
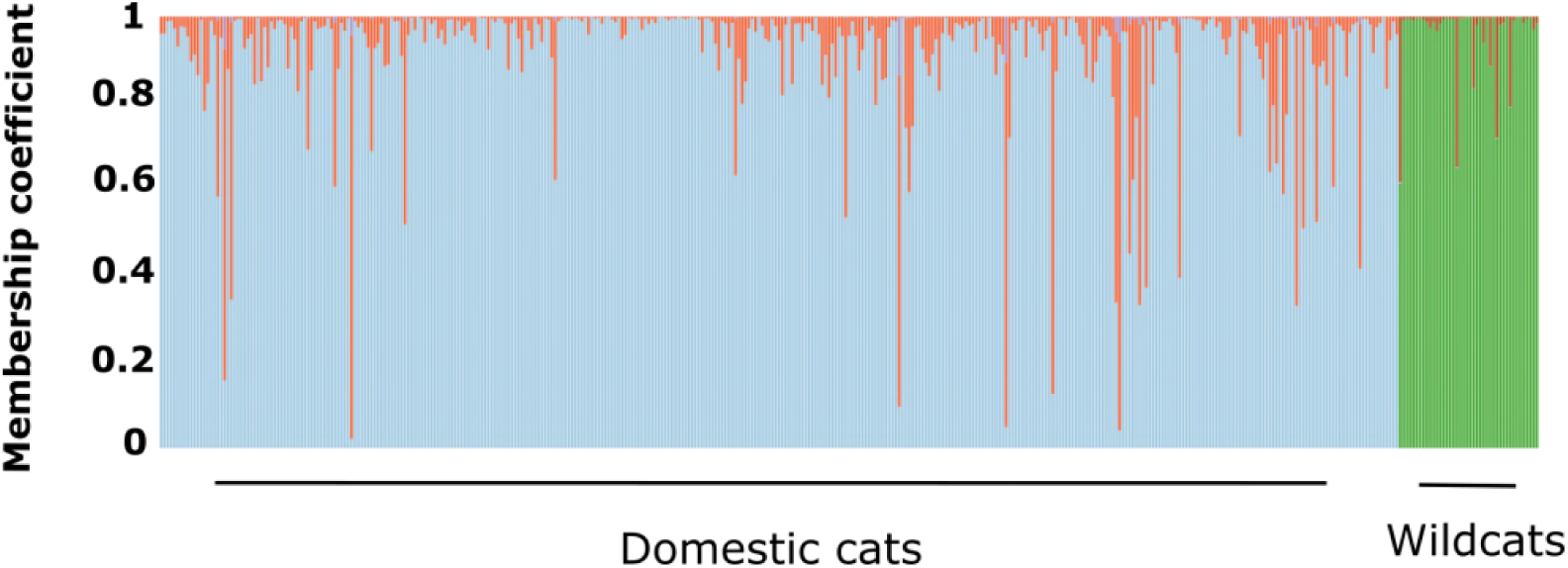
Results from the *snapclust* analysis for the northeastern population. Each vertical bar represents an individual. The colors represent the different hybrid categories included in the analysis: domestic cats in blue, wildcats in green, F1 in purple, F1xdomestic cats in orange, F1xwildcats in brown. The proportion of the bar in a given color represents the probability for the individual to belong to the corresponding category.

